# Amygdala neuronal ensembles dynamically encode behavioral states

**DOI:** 10.1101/425736

**Authors:** Jan Gründemann, Yael Bitterman, Tingjia Lu, Sabine Krabbe, Benjamin F. Grewe, Mark J. Schnitzer, Andreas Lüthi

**Author notes:** these authors contributed equally (Jan Gründemann, Yael Bitterman). present address: University of Basel, Department of Biomedicine, Klingelbergstrasse 50-70, Basel, Switzerland Contact (Jan Gründemann).

## Abstract

Internal states, including affective or homeostatic states, are important behavioral motivators. The amygdala is a key brain region involved in the regulation of motivated behaviors, yet how distinct internal states are represented in amygdala circuits is unknown. Here, by imaging somatic neural calcium dynamics in freely moving mice, we identify changes in the relative activity levels of two major, non-overlapping populations of principal neurons in the basal nucleus of the amygdala (BA) that predict switches between exploratory and non-exploratory (defensive, anxiety-like) behavioral states across different environments. Moreover, the amygdala widely broadcasts internal state information via several output pathways to larger brain networks, and sensory responses in BA occur independently of behavioral state encoding. Thus, the brain processes external stimuli and internal states in an orthogonal manner, which may facilitate rapid and flexible selection of appropriate, state-dependent behavioral responses.

## Introduction

Internal state-dependent adaptations of perception and selection of appropriate behavioral outputs are essential for an animal’s survival^1^. Changes in internal states induced by running or whisking^2,3^ have been linked to distinct states of thalamocortical circuitry and sensory coding^4–6^. However, to date it has remained elusive where and how complex affective and homeostatic internal state changes are encoded as neural state changes, such as by specific neural ensembles that change their patterns of activity across different behaviors. Prior studies identified state-related functions of subcortical^7^ brain areas and neural circuits^8^ via electrical and optogenetic stimulation, which can induce rapid shifts in affective or defensive behaviors, foraging drives, feeding behavior and systemic hormonal levels^9–16^. Homeostatic (hunger and thirst) or affective internal states induced by aversive or appetitive experience are each associated with perceptual, autonomic and hormonal responses, which lead to distinct behavioral outputs, e.g. anxiety-like, defensive behaviors upon predator exposure^17–23^. This suggests that internal states are represented by specific patterns of neuronal activity across large brain networks^24^.

Nevertheless, how ensembles of identified neurons can represent internal states via their activity patterns has remained unknown^2,25,26^, partly due to the lack of suitable techniques for tracking of behavioral and neuronal activity. By using recently developed methods for deep brain optical Ca^2+^ imaging^27,28^ in identified neuron types of freely moving animals, here we identify neural ensemble representations of internal states and illustrate their generality across multiple different behavioral contexts and paradigms.

The basolateral amygdala (BLA) and specifically its basal nucleus (BA), is a brain hub for regulating affective, homeostatic and social behaviors. Notably, the BA is functionally linked to motor pathways^29^ that drive specific behavioral outputs, thalamic and cortical sensory areas that process outcome-predicting stimuli^30,31^, and brain centers that sense and regulate behaviorally-relevant hormones and neuromodulators^32^. Previously, amygdala function has mainly been studied in the framework of appetitive and aversive Pavlovian conditioning^33–36^, yielding specific circuit elements that underlie associative plasticity at the single cell^37–39^ and neural ensemble level^28^. Thus, while past work has well characterized amygdalar activity that is learned and temporally locked to experimentally controlled events, such as the conditioned and unconditioned stimuli used for Pavlovian conditioning, we only have a rudimentary understanding of how amygdalar neural activity relates to self-paced, state-driven behaviors, including foraging, risk assessment or place avoidance^40,41^. These behaviors are strongly driven by internal states, such as exploratory drive or anxiety^42^, and thus may therefore serve as their external manifestation and read-out; yet how ensembles of identified BA neurons encode these states and their relationships to learned stimulus-outcome representations remains unknown.

To determine how amygdalar neural ensembles^24^ represent internal states, we performed deep brain Ca^2+^ imaging studies of large populations of BA principal neurons (PNs) using a head-mounted miniaturized microscope^27,28^ in mice that engaged in a series of behavioral paradigms, including those traditionally used to manipulate anxiety levels as well as Pavlovian fear conditioning. By longitudinally tracking large ensembles of individual neurons across several days and paradigms (Fig. S1) and using three complementary approaches to analyze ensemble neural dynamics, we identified a prominent, trackable neural population signature of internal states in the BA. This signature is reminiscent of neural states and is invariant across multiple days and behavioral paradigms and can be described as two opposing activity patterns arising in two functionally distinct neural ensembles. Further, these two different population states are broadcast to multiple downstream brain areas and do not correlate with sensory-driven BA activity, indicating that behaviorally relevant internal states and sensory inputs seem to be encoded in an orthogonal manner. The neural population signature has slow, seconds-scale dynamics, manifests as fluctuations in population level activity^43^ throughout a behavioral session, and arises during diverse exploratory and non-exploratory, defensive behaviors on consecutive days, thereby enabling accurate predictions of the animal’s behavioral output.

### Amygdala activity during open field exploration

To investigate general principles of BA population coding, we used a miniature fluorescence microscope (Fig. 1A, Fig. S1A-D) to track the relative changes in Ca^2+^ fluorescence in large populations of BA PNs (CaMKII-positive as well as projection-specific PN populations, Fig. S7) in freely moving mice within three different, consecutive behavioral assays; an open field test, OF; elevated plus maze test, EPM; auditory fear conditioning, FC; (Fig. 1B; Fig. 5A, Fig. S1). We chose these assays because they prompt mice to exhibit distinct modes of behaviors that are likely outward manifestations of different internal states.

**Fig. 1.**
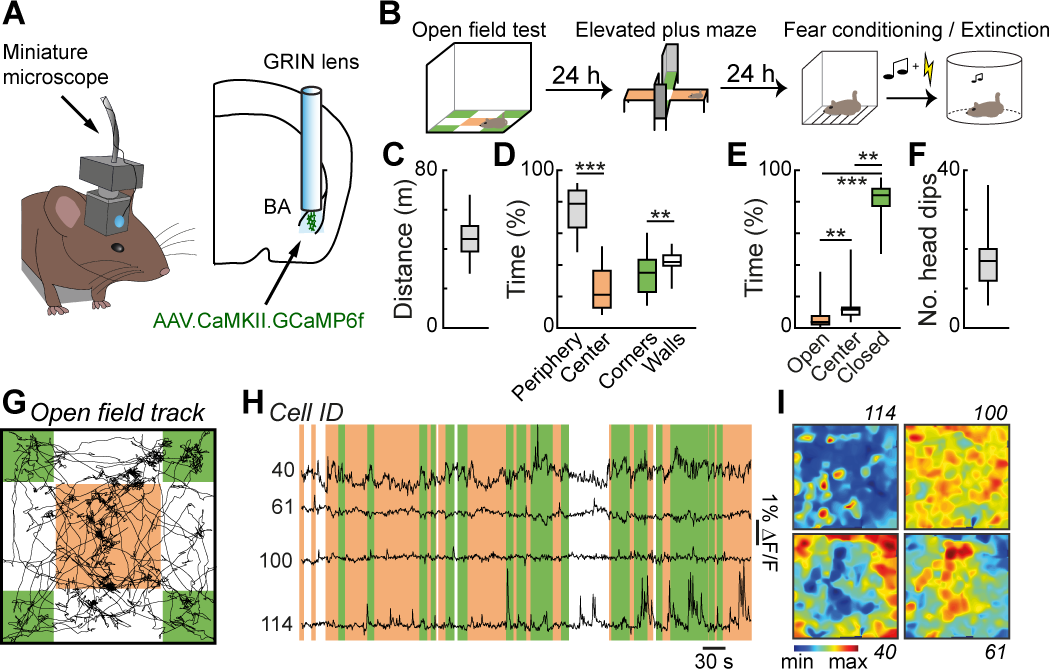
Deep brain imaging of BA activity in freely exploring mice. **A)** Scheme of gradient index lens implantation and virus expression strategy. **B**) Scheme of seven-day behavioral paradigm including consecutive open field test, elevated plus maze test as well as five day fear conditioning and fear extinction paradigms (see also Fig. 5). **C**) Distance travelled in open field (N = 25 mice). **D**) Time spent in center vs. periphery and in subareas of the periphery (walls vs. corners). ** P < 0.01, *** P < 0.001, Wilcoxon matched-pairs signed rank test (N = 25 mice). For definition of locations see G. **E**) Elevated plus maze: Time-spent in open and closed arms as well as center. **F**) Average number of head dips on the elevated plus maze (N = 25 mice). **G**) Example open field track of an individual mouse. Context size: 40×40 cm. **H**) Example Ca^2+^ signals of four simultaneously recorded individual cells during open field exploration. Colors indicate mouse location: orange-center, green-corner. **I**) Mean Ca^2+^ signal across the open field arena for the four example cells shown in H demonstrating non-specific (cell 114 and 100) and area-modulated activity patterns (cell 40 and 61). Color bar indicates normalized z-score. Box and whisker plots indicate median, interquartile range and the minimum to maximum values of the data distribution.

During the open field test, mice generally behaved as previously described^44–47^ and spent most time in the periphery of the OF (Fig. 1D, periphery: 76 ± 3%, center: 24 ± 3%, P < 0.0001, Wilcoxon matched-pairs signed rank test, N = 25 mice). Nevertheless, mice typically exhibited pronounced exploratory behavior in the OF (Fig. 1G, Fig. S2G). They covered large distances (45 ± 2 m in ca. 10 min, N = 25 mice) (Fig. 1C) and ventured out of the corners along the walls (time in corners: 34 ± 2%, time at walls: 42 ± 1%, P < 0.01, Wilcoxon matched-pairs signed rank test, N = 25 mice) and into the center (Fig. 1D). We tracked the somatic Ca^2+^ activity of BA CaMKII-positive PNs (133 ± 6 neurons / animal, N = 9 mice) throughout OF exploration (Fig. 1A, Fig. S1). BA neurons exhibited diverse activity patterns that included discrete Ca^2+^ transient events as well as slow changes in Ca^2+^-related fluorescence (Fig. 1H). The average spatial response patterns of individual BA neurons in the OF arena ranged from seemingly non-specific to area-biased activity that preferentially occurred when the mouse was either in the arena corners (corner-modulated cells) or center (center-modulated cells) (Fig. 1I). We quantified the area-bias of the neuronal responses based on the differential Ca^2+^ activity between corners and centers (area score, see Methods). We defined neurons with scores exceeding a selected threshold as area-modulated (total area modulated: 28 ± 5%) (Fig. 2A, B). The area score distribution based on differential activity in corners vs. centers was significantly wider than that of area scores based on neutral divisions of the open field (e.g. left vs. right, top vs. bottom) or datasets in which we temporally shifted the cellular activity traces relative to the behavioral time course (Fig. S2A, see Methods). Accordingly, a substantial fraction of corner vs. center-based area scores was outside of the 95% confidence interval of the neutral left vs. right (17%) or top vs. bottom (21%) distributions, as well as the temporally shifted controls (21 ± 3% for 100 random temporal shifts), indicating that the extent of corner versus center coding in BA ensembles is significantly greater than that expected from the random fluctuations in the cells’ activity patterns. Across the population, there were comparable proportions of cells that were preferently active in the corners or in the center (corners: 13 ± 3% of BA PNs / animal; center: 15 ± 3%; P = 0.641, Wilcoxon matched-pairs signed rank test; N = 9 mice) (Fig. 2C). Corner-modulated and center-modulated neurons were spatially intermingled in the BA with no apparent local clustering (Fig. 2D, E, Fig. S2B).

**Fig. 2.**
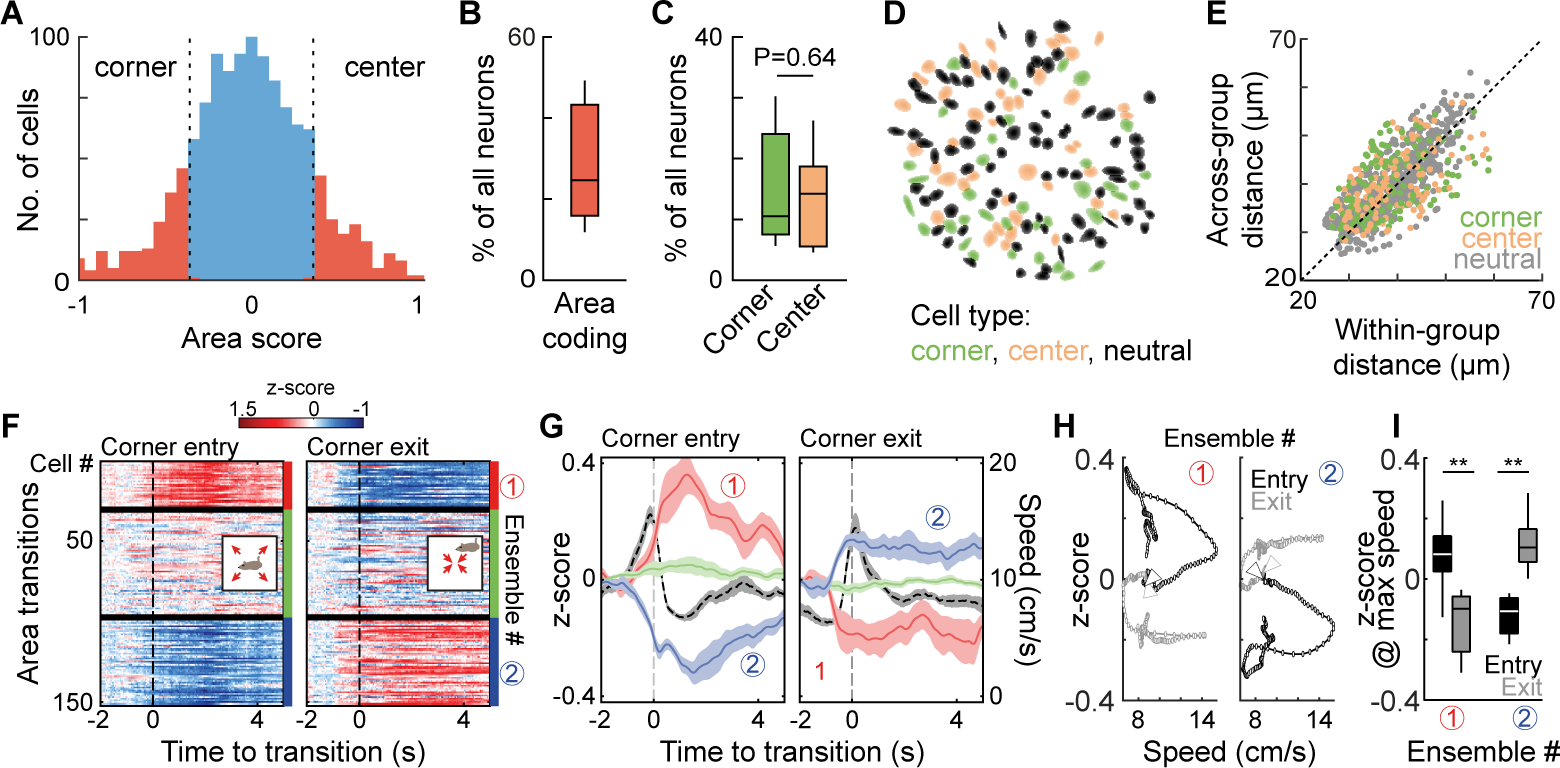
Large, intermingled populations of basal amygdala neurons are spatially modulated. **A**) Histogram of area scores of CaMKII-positive amygdala projection neurons (n = 1201 neurons from 9 mice). Cut-off scores for corner-and center-modulated cells were set at ± 1 standard deviation (vertical dashed lines). **B**) Proportion of area coding neurons. **C**) Proportion of corner-and center-modulated neurons is similar (P = 0.64, Wilcoxon matched-pairs signed rank test). **D**) Spatial distribution of corner-(green) and center-modulated (red) cells in the BA of one example mouse. Black: neutral cells. **E**) Mean within and across-group distance of individual corner-and center-modulated cells (each point represents one cell, see also Fig. S2B). **F**) Mean neuronal responses of functionally clustered groups of individual neurons upon corner entry (middle) and corner exit (right) transitions (n = 152 simultaneously imaged neurons from one mouse, number of entries: 24, number of exits: 21). Zero marks corner entry or corner exit times, respectively (see methods) and responses were baselined using the first 0.5 s response (starting 2 s before transition). Cells were ordered according to K-means clustering on the average corner entry response of each cell and cell IDs were kept identical for average corner exit responses. **G**) Average corner entry (left) and exit (right) response of corner entry activated (red, ensemble 1), neutral (green) or inhibited (blue, ensemble 2) clusters across animals. Black traces indicate the average speed profile upon corner entry or exit. Lines indicate average response across animals ± s.e.m‥ As in the example animal shown in G, cluster IDs were based on corner entry responses and kept for corner exits. **H**) Ca^2+^ activity in relation to corner entry (black) and corner exit (grey) transition speed for the two ensembles (average response profile for N = 9 animals) reveals that the relationship between instantaneous speed and cellular activity depends on the spatial destination and is not solely speed-driven. Triangles mark the start of the transition. **I**) Average z-score at maximum transition velocity (see H, Ensemble 1: P = 0.008, Ensemble 2: P = 0.004, Wilcoxon matched-pairs signed rank test). Box and whisker plots indicate median, interquartile range and the minimum to maximum values of the data distribution. All data is generated from N = 9 animals.

To examine neuronal response dynamics during an animal’s transition from the OF center into a corner, and vice versa, we averaged neuronal Ca^2+^ responses aligned on crossings of the corner boundary according to the directionality of the transition. This analysis revealed that a large group of BA neurons was either activated (27 ± 4%, hereon termed ensemble 1) or inhibited (29 ± 4%, hereon termed ensemble 2, P = 0.641, Wilcoxon matched-pairs signed rank test, see Methods for details on cluster based analysis of relative changes in Ca^2+^ fluorescence) upon spatial transitions into or out of the corners (Fig. 2F, G). Interestingly, the sign of the response of the respective BA ensemble was inverted when the animal exited the corner area, notwithstanding similar speed profiles for both transitions (Fig. 2G, H, Fig. S2J). Generally, ensemble 1 cells activated upon corner-entry and overlapped with corner-modulated cells, whereas ensemble 2 cells activated at corner-exits and corresponded to center-modulated cells (Fig. 2F, Fig. S2C-F). The activity patterns of corner‐ and center-modulated cells were linked to the spatial location or the transition therein and did not depend on differences in average corner and center movement (Fig. S2G), general speed correlations on the single cell level (Fig. S2H), or correlations of neuronal activity with instantaneous changes in speed (Fig. 2G-I, Fig. S2I, J). Altogether, although BA neurons can be speed modulated, speed modulation alone cannot explain the prominent area coding.

Given the strong area-modulation of the neuronal responses, we next trained a support vector machine classifier (see Methods) to test if corner vs. center locations of the animal could be accurately predicted solely based on BA population activity. Indeed, the classifier reached high decoding accuracies of 86 ± 2%, whilst decoder performance dropped if the decoder was trained and tested on temporally shuffled neuronal data (46 ± 4%, P = 0.004, Wilcoxon matched-pairs signed rank test) (Fig. S2K) or temporally shifted to control for local structures in the behavioral and neuronal data (Fig. S2L, see also Methods).

Finally, we tested the extent to which corner‐ and center-coding was stable across days by using an open field re-exposure paradigm. Area scores of individual neurons were significantly correlated (R = 0.53, p < 0.001) across days, indicating the stability of BA area coding (Fig. S3).

Taken together, this data shows that, during the OF exploration, a large fraction of BA neurons conveys area-modulated information about an animal’s corner or center location. Notably, such areas are generally thought to reflect different internal states with regard to the animal’s anxiety or stress levels^45,48–50^.

### Amygdala encoding of elevated plus maze exploration

To investigate if there are general principles of BA population coding that apply to exploratory behavior across different contexts, we imaged the same BA PNs during EPM exploration (Fig. 1B, 3B, Fig. S1E). In keeping with the OF test, mice spent more time in the closed arms of the EPM (closed: 81 ± 2%). Typically, one of the closed arms, the preferred home base arm, was substantially more frequented (Fig. S4A-D). Similar to our observations in the OF test, mice also exhibited exploratory behavior on the EPM. They ventured out of the closed arms into the center (relative time spent: 13 ± 2%) or onto the open arms (6 ± 1%, P < 0.0001, Friedman test) (Fig. 1E). Furthermore, the animals performed so-called head dips, characterized by brief periods of visual investigation over the edge of the EPM open arms (Fig. 1F).

Next, we asked whether the area-modulated responses of BA PNs which we observed in the OF would generalize to the EPM. Specifically, we hypothesized that enclosed, potentially safe areas are similarly represented between the two paradigms, such that corner-modulated cells might be more active during an animal’s stay in the closed arm of the EPM, whereas center-modulated cells might show higher activity levels during open arm exploration. Surprisingly, the average activity of corner-modulated cells was reduced in the preferred closed arm (mean z-score: −0.22 ± 0.03, N = 9 animals) (Fig. 3B, E), whereas center-modulated cells exhibited increased activity in the preferred closed arm (mean z-score:0.15 ± 0.04, P < 0.01, Wilcoxon matched-pairs signed rank test) (Fig. 3C, E). Once the animal ventured out of its preferred closed arm to explore the EPM, the average activity of corner‐ and center-modulated cells reversed such that corner-modulated cells increased their Ca^2+^ fluorescence in the center (Δz-score: 0.246 ± 0.04), less frequented closed arm (Δz-score: 0.262 ± 0.04) or open arms (Δz-score:0.222 ± 0.03), and vice-versa center-modulated cells (Δz-score for center = −0.177 ± 0.04, closed-avoided = −0.183 ± 0.05, open = −0.156 ± 0.05; all P < 0.01, Wilcoxon matched-pairs signed rank test) (Fig. 3D-F). This suggests that the activity of corner-modulated cells correlates with exploration of the non-preferred closed and open arms of the EPM. Consistent with this notion, the animals’ behavior in the OF corners was not passive, but rather characterized by constant activity, exploration and rearing (Fig. S2G and S4E).

**Fig. 3.**
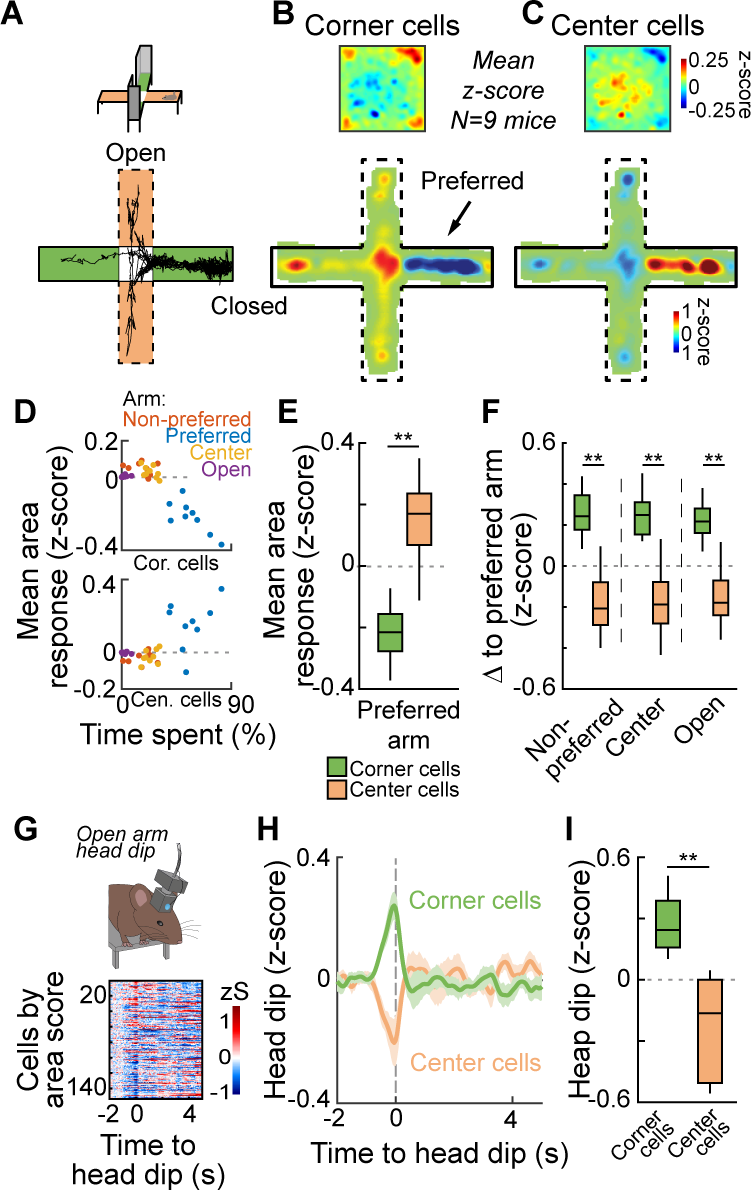
Corner-and center-modulated cells encode exploratory behavior. **A)** Elevated plus maze behavior (EPM) of the example animal shown in Figure 1. Green areas represent closed arms and orange areas represent open arms of the elevated plus maze. **B**) Average spatial Ca^2+^ activity map of open field corner-and center-modulated cells in the open field (top) and on the EPM (bottom) across animals. **C**) Average spatial Ca^2+^ activity map of center-and corner-modulated cells in the open field (top) and on the EPM (bottom) across animals. Animals typically have a preferred closed arm and EPM maps were rotated such that the most frequented closed arm is aligned on the right (arrow in C). **D**) Mean area responses (N = 9 animals) in the preferred closed arm (blue), non-preferred closed arm (red), EPM center (yellow) and open arms (lilac) for corner (top) and center (bottom) in relation to the relative time spent at each location. The mean spatial activity in relation to the time spent at the location are inverted for corner and center cells. **E**) Average preferred arm response of corner and center cells (P < 0.01, Wilcoxon matched-pairs signed rank test). **F**) Response difference of non-preferred arms, open arms as well as center to the preferred, home base arm for corner-and center-modulated cells (P < 0.01 for all comparisons, Wilcoxon matched-pairs signed rank test, E and F based on non-time-normalized data). **G**) Illustration of head dip behavior on EPM open arms (top) and example BA responses to head dips for 152 simultaneously recorded cells of one animal (bottom). Cell IDs sorted by area score (see also Fig. 2A). **H**) Average response of corner-and center-modulated cells upon EPM head dips. **I**) Quantification of peak response shown in h (P < 0.01, Wilcoxon matched-pairs signed rank test). Box and whisker plots indicate median, interquartile range and the minimum to maximum values of the data distribution. All data is generated from N = 9 animals.

To test this idea further, we analyzed activity of corner-and center-modulated cells during open arm head dips, a classic exploratory, low anxiety behavior^51^. On average, the animals performed 18 ± 2 head dips per EPM session. As expected, during head dips, corner-modulated cells were strongly activated (z-score: 0.28 ± 0.04) whereas center-modulated cells were inhibited (z-score: −0.23 ± 0.08, P < 0.01, Wilcoxon matched-pairs signed rank test) (Fig. 3G-I). These data demonstrate that the opposing activation pattern of corner-and center-modulated cell ensembles is preserved during head dips, an alternative, stationary exploratory behavior. Together, these data indicate that two large, non-overlapping functional ensembles of BA PNs signal fundamentally distinct states characterized by the presence or absence of exploratory behavior across different paradigms.

### Consistent encoding of behavioral states in amygdala ensembles

Next, we asked how behavioral states and the transitions between them are encoded on the neuronal population level. We extended the analysis to follow the modulation of BA ensemble activity across different states and behavioral paradigms. We classified the neuronal ensembles based on their responses during transitions into the corners of the OF into activated (ensemble 1, or corner-entry activated, 27 ± 4% of all neurons), neutral (transition neutral, 44 ± 3%) or inhibited (ensemble 2, or corner-exit activated, 29 ± 4%) neurons (Fig. 4A, see also Fig. 2G). The activated and inhibited ensembles showed on average inverted relative changes in Ca^2+^ fluorescence upon corner exit (Fig. 4A-C, Fig. 2G). We next analysed the activity of these three ensembles aligned on different behavioral transitions using the same cluster partition and cell IDs. Indeed, consistent with the idea that ensemble 1 neurons encode a general state associated with exploratory behavior across animals, ensemble 1 cells were not only activated upon OF corner entries, but also upon open arm head dip behavior (Fig. 4A-C, see also Fig. 3G-I). In contrast, ensemble 2 neurons exhibited the inverse pattern of activity modulation (Fig. 4A-C). Notably, when extending this analysis to freezing behavior (see Fig. 5A), a well-studied, distinctive defensive behavior induced by auditory fear conditioning, we found that ensemble 1 and ensemble 2 neurons showed opposite activity changes when animals transitioned into a high fear, freezing state and upon termination of freezing (Fig. 4A, for details see Methods).

**Fig. 4.**
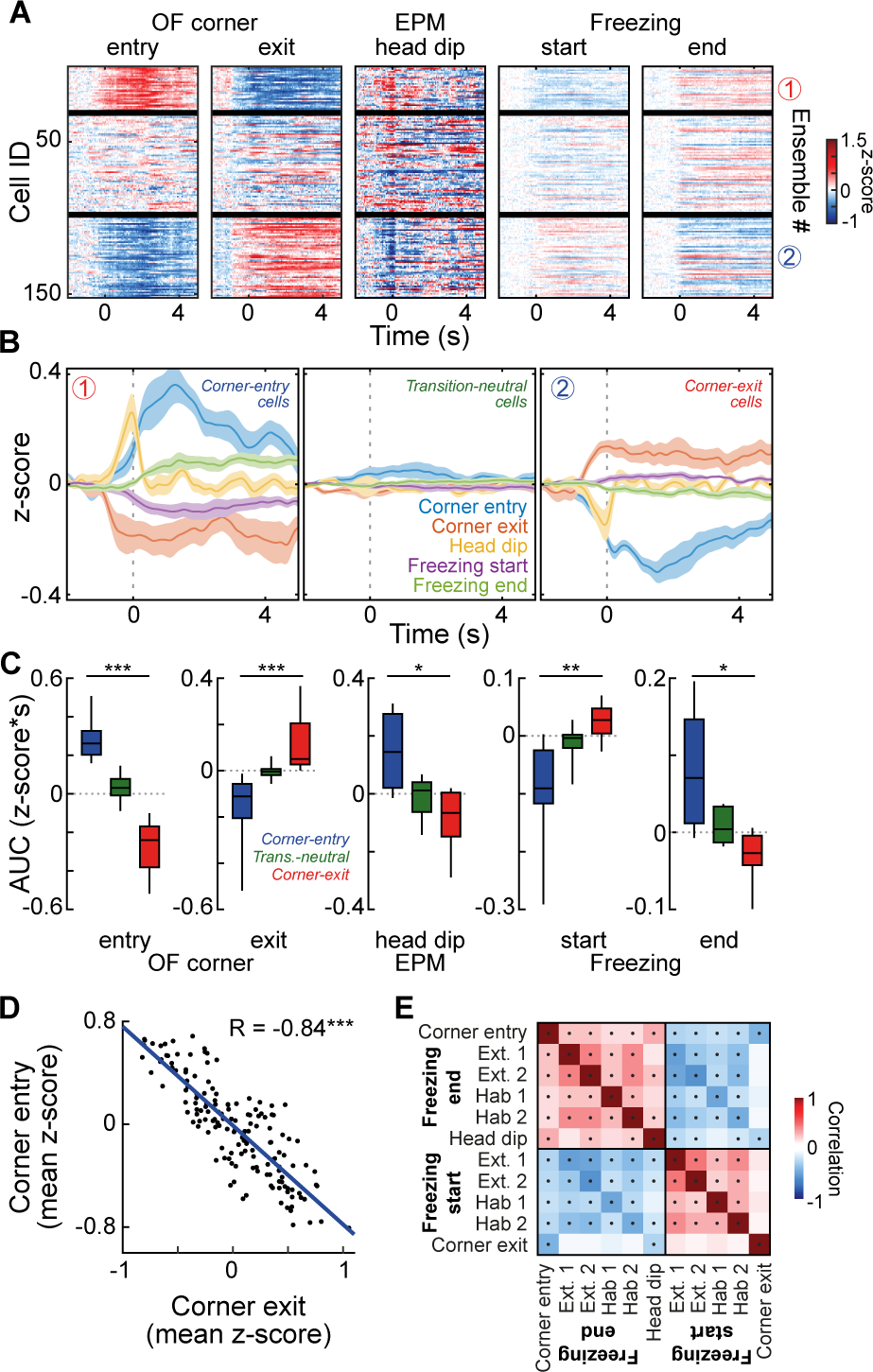
State-coding of BA neuronal ensembles across behavioral paradigms. **A)** Average response patterns of open field corner entry, corner exit EPM head dips and freezing start as well as freezing end for cells clustered based on OF corner entry responses (top). **B**) Average cluster response across animals (N = 9). **C**) Quantification of the average cluster responses aligned to open field corner entry (z-score: Corner-entry cells (Ensemble 1) = 0.28 ± 0.04, Transition-neutral cells = 0.03 ± 0.02, Corner-exit cells (Ensemble 2) = −0.27 ± 0.05, P < 0.001 for Corner-entry cells vs. Transition-neutral cells), to corner exit (Corner-entry cells = −0.16 ± 0.05, Transition-neutral cells = −0.003 ± 0.01, Corner-exit cells = 0.11 ± 0.04, P < 0.001 for Corner-entry cells vs. Transition-neutral cells), to EPM head dips (Corner-entry cells = 0.15 ± 0.04, Transition-neutral cells = −0.01 ± 0.02, Corner-exit cells = −0.09 ± 0.03, P = 0.014 for Corner-entry cells vs. Corner-exit cells) as well as to freezing start (Corner-entry cells = −0.09 ± 0.03, Transition-neutral cells = −0.01 ± 0.01, Corner-exit cells = 0.03 ± 0.01, P = 0.003) and to freezing end (Corner-entry cells = 0.08 ± 0.02, Transition-neutral cells =0.01 ± 0.01, Corner-exit cells = −0.03 ± 0.01, P = 0.020, all Friedman test with Dunn’s multiple comparison test). **D**) Single cell correlation of corner entry and corner exit responses of one example animal (N = 152 cells, R = −0.84, P < 0.001). **E**) Average correlation matrix of single cell responses across different behavioral paradigms (dots indicate a significant difference of the correlation from 0 across animals, one sample t-test). Box and whisker plots indicate median, interquartile range and the minimum to maximum values of the data distribution. All data is generated from N = 9 animals.

**Fig. 5.**
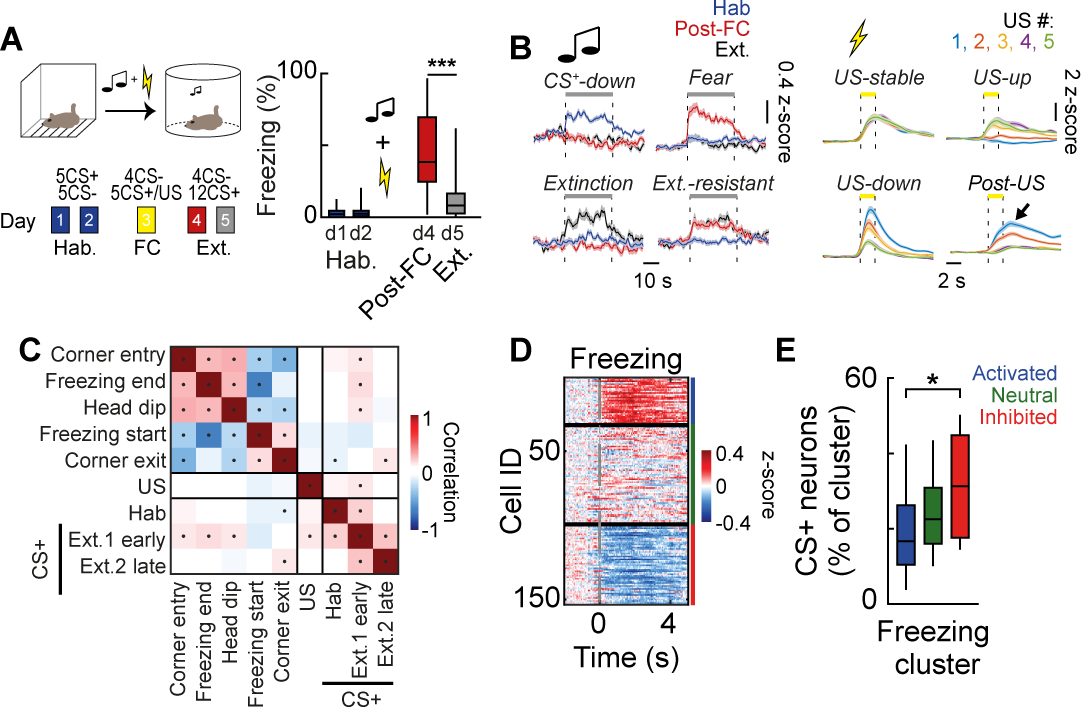
Orthogonal representation of sensory stimuli and behavioral states. **A)** Auditory fear conditioning paradigm and average CS+ freezing (Post FC freezing: 44 ± 6 %, post extinction freezing: 13 ± 3 %, P < 0.001, Wilcoxon matched-pairs signed rank test). **B**) Left: Average CS+ responses of CS-down, fear, extinction (Ext.) and extinction-resistant (Ext.-resistant) neurons. Right: Average US responses of US-stable, US-up, US-down, and Post-US excited neurons. **C**) Single cell correlation matrix of behavioral as well as CS and US responses across animals (dots indicate significant correlations across animals). **D**) Clustered freezing responses of 152 neurons of one example animal. Color bar indicates z-score. **E**) Average proportion of CS+ and freezing cluster-responsive neurons out of the total population (CS+ / freezing activated = 18 ± 4 %, CS+ / freezing neutral = 24 ± 3 %, CS+ / freezing inhibited = 32 ± 5 %, P(activated vs. inhibited) = 0.029, Friedman test (P = 0.03, F = 6.89) with post hoc Dunn’s multiple comparison test). Box and whisker plots indicate median, interquartile range and the minimum to maximum values of the data distribution. All data is generated from N = 9 animals.

Since ensemble responses were averaged across many neurons (44 ± 3 cells / cluster), we verified that similar results are obtained for single neuron responses using an alternative response correlation measure. Single neuron responses were aligned to the same behavioral transitions used for the ensemble analysis and pair-wise correlation was calculated between responses to a pair of behaviors across the entire neuronal population (N = 1201). As expected, single neuron responses to corner entry and corner exit were highly negatively correlated (R = −0.84, P < 0.001) (Fig. 4D). In further agreement with the ensemble analysis, correlations between single neuron responses to corner entry, freezing end and head dips were highly positive, whereas the correlations to freezing start and corner exit behaviors were significantly negative (Fig. 4E). These correlations were independent of the behavioral session (i.e. freezing during habituation or extinction 1 and 2) (Fig. 4E). Together, these data reveal the population level encoding of behavioral states by large, antagonistic ensembles of BA neurons, which switch their activity patterns upon opposing exploratory and anxiety-like or defensive behavioral states.

### Orthogonal behavioral and sensory coding in BA neuronal ensembles

Next, we tested how the population coding of behavioral states in BA neurons relates to the population coding of sensory stimuli during the acquisition and extinction of classical Pavlovian fear conditioning. During conditioning, we paired a previously neutral pure tone (conditioned stimulus, CS+, 75 dB pure tone) with a mild electrical foot shock (unconditioned stimulus, US, 2 s, 0.65 mA AC, see Methods). After conditioning, the animals showed increased freezing responses towards the CS+, which extinguished upon repeated re-exposure to the CS+ (Fig. 5A). BA neurons were ca. two times more likely to respond to the US (57 ± 3% of all neurons including excitatory and inhibitory responses, N = 9 animals) (Fig. S5A-C) than to the CSs (30 ± 3% of all neurons, P = 0.004, N = 9 animals; Wilcoxon matched-pairs signed rank test, for selection criteria see Methods) and were more likely to be activated by the CS+ than the CS−(CS+ 25 ± 3%, CS− 8 ± 1%, P = 0.004, Wilcoxon matched-pairs signed rank test) (Fig. S5D).

We classified neurons according to the evolution of their CS+ responses upon fear conditioning and extinction, using a supervised clustering approach (Fig. 5A, Fig. S5A, see Methods). Consistent with previous findings^28,38,52–54^, we found functionally distinct CS plasticity subtypes of BA PNs that were characterized by differential changes of CS+ responses during fear conditioning and extinction, for example, CS-down neurons, fear neurons, extinction neurons or extinction-resistant neurons (Fig. 5A, Fig. S5A, E). Interestingly, none of these CS plasticity subtypes were correlated with either of the behavioral state-related neuronal ensembles (Fig. S6A), indicating that experience-dependent CS representations are maintained across different behavioral states.

US responses were dynamic upon fear conditioning (Fig. 5B). Across the five CS-US pairings, a supervised cluster analysis (Fig. S5B) identified different subtypes of US-excited (32 ± 2% of all neurons, US-stable, US-down, US-up as well as post-US neurons) and US-inhibited neurons (25 ± 2% of all neurons, compared to US-excited: P = 0.129, Wilcoxon matched-pairs signed rank test, stable US-inhibited and plastic US-inhibited down neurons) (Fig. S5F, G). Despite the detailed classification of highly diverse and dynamic response patterns to the CS and US across the entire population of BA PNs, no significant overlap between individual CS and US plasticity subtypes was observed (Fig. S5H; Fig. S6A, C, see also Fig. 4). Notably, a population level analysis revealed that US responses per se were not correlated with any of the tested behavioral states (Fig. 5C, Fig. S6B), indicating that US representations are not specifically associated with a particular behavioral state. Overall, these results demonstrate that, on the level of BA neuronal populations, sensory representations of the US are uncorrelated with and are encoded orthogonally to behavioral state-related population activity.

A more detailed analysis of CS+ responses across behavioral paradigms revealed that before fear conditioning and after fear extinction, the population level CS+ response was not correlated with exploratory or freezing behaviors, which is consistent with the neutrality of the CS+ response at these time points. Strikingly though, after fear conditioning, the CS+ population response was weakly correlated with exploratory, low anxiety behaviors (corner entries, freezing end, head dips, Fig. 5C). Consistent with this observation, we found that after fear conditioning, more CS+ responsive neurons overlapped with freezing start-inhibited neurons (32 ± 5%), which are typically activated by exploratory behaviors (similar to ensemble 1, see Fig. 4), than with freezing start activated neurons (18 ± 4%, similar to ensemble 2, P = 0.029, Friedman test (P = 0.03, F = 6.89) with *post hoc* Dunn’s multiple comparison test, Fig. 5D, E, for US see Fig. S6B). A stronger CS+ representation in freezing-inhibited neurons, separating state-activated ensembles from sensory coding neurons, may respresent a possible circuit mechanism to enhance the signal-to-noise ratio for salient sensory stimuli such as a CS after conditioning.

Finally, given previous reports indicating that defined BA output pathways are involved in specific behaviors^37–39,55,56^, we tested the hypothesis that the BA would transmit distinct behavioral state-or sensory-related representations to select downstream targets including the ventral hippocampus (vHC), the medial prefrontal cortex (mPFC) or the nucleus accumbens (NAc). Although some specialization could be observed for individual pathways (typically CS/US coding-related^57^), the population correlations with behavioral states was largely maintained across output pathways (Fig. S7), suggesting that the BA broadcasts state-related signals to larger brain networks.

## Discussion

Our study reveals a novel coding principle of internal state representations in large, functional ensembles of BA neurons (Fig. S8). Using a longitudinal time-lapse imaging approach across multiple paradigms in freely moving animals, we demonstrate large-scale opposing activity dynamics in two functionally distinct, anatomically intermingled ensembles of amygdala neurons that code for distinct behavioral state transitions. Importantly, state-coding in amygdala ensembles was consistent across different behavioral paradigms and changes in state-coding reflected transitions from exploratory behavior to non-exploratory, defensive, anxiety-like states and back. These neural state signatures were not exclusively driven by changes in movement parameters of the animal, but were direction selective and represented similarly in stationary (e.g. head-dips) and non-stationary exploratory behaviors. Our findings in freely moving animals extend the concepts of cortical state coding classically related to movement-and arousal/attention-related processes in brain state^43,58–63^ towards dynamic exploratory and affective state representations by large neural ensembles in subcortical circuits that may include diverse patterns of physiological, sensory and locomotion processes.

Consistent state-dependent changes in the activity of single cells suggest a fixed state ensemble membership of individual neurons and a homogenous, state-dependent, regulation of individual neuron activity within a given state ensemble. Strikingly, changes in population activity were characterized by slow (seconds), continuous dynamics in defined ensembles and aligned with behavioral transitions, reminiscent of a potential amygdala attractor network state, which has been previously suggested to underly internal states^64^. Slow changes in amygdala ensemble activity that emerged as a basic structure on the population level were predictive of changes in exploratory behaviors across days and paradigms (Fig. S9). Such slow regulation of antagonistic BA ensemble activity might involve state-dependent changes in input activity, neuromodulation and local circuit mechanisms^65–68^ and can now be tested for generality in other freely moving behavioral paradigms.

Notably, despite being encoded by amygdala neuronal populations^28^, sensory responses were correlated only partially (CS), or not at all (US), with the coding of behavioral states. This suggests that, similarly to cortex^58,59^, fast sensory dynamics and slow state dynamics can be orthogonally separated at the level of amygdala networks, which would allow for a continuous population level state-encoding despite flexible single cell representations of sensory inputs^69^. Additionally, biased representations of emotionally salient stimuli in freezing-inhibited neurons might substantially increase signal to noise ratio and enhance the animal’s ability to rapidly and reliably select appropriate behavioral reactions when facing danger. Beyond freezing, the concurrent signalling of both, the conditioned stimuli (CS) and the learned, conditioned responses (CR) in the BA^70^ would not only allow for a flexible, state-driven modulation of CS representations, but also enable appropriate CR selection in a state-dependent manner. Importantly, correlations of responses to the behavioral state transitions were by-and-large conserved across major BA output pathways including projections to the vHC, the NAc and the mPFC, despite individual differences between CS/US coding in these projections. While we cannot rule out projection-target differences of state coding in relation to local gamma entrainment^68^, broadcasting of amygdala state signals to larger brain networks including cortical and subcortical areas is likely to play an important role for state-dependent regulation of sensory perception, motivation^71^, foraging^41^, spatial preference^72^, memory formation^73^, as well as motor output^29^ and autonomic regulation^32^, with a potential to broadly modulate brain-state^4,59^ and internal drive and generally modify affective states and behaviors.

## Acknowledgements

We thank Georg Keller and Daniela Gerosa Erni for virus production and V. Jayaraman, R. Kerr, D. Kim, L. Looger, K. Svoboda and the HHMI Janelia GENIE Project for making GCaMP6 available as well as Alla Karpova and David Schaffer who gifted the rAAV2-retro helper (Addgene plasmid # 81070V). We thank Paul Argast and Peter Buchmann for workshop and engineering support. Research was supported by the Swiss National Science Foundation (Core Grant to A.L., Ambizione Fellowship to J.G., SNF Professorship to J.G., Sinergia to B.F.G.), EMBO and Marie Curie Actions (J.G.), NARSAD (Young Investigator Fellowships to S.K. and B.F.G.), Swiss Data Science Center (B.F.G.), DARPA (M.J.S.), NIH (M.J.S), ERC (Advanced Grant, A.L.) and the Novartis Research Foundation.

## Author contribution

J.G. Y.B., and A.L. designed experiments and analysis. J.G., T.L. and S.K. performed experiments. J.G., B.F.G. and M.J.S. established the Ca^2+^ imaging protocol, experiments and image data analysis. Y.B. and J.G. analysed data. J.G., Y.B. and A.L. wrote the paper and all authors edited the paper and commented on the manuscript.

## Methods

### Animals and viruses

Male C57BL/6J mice (C57BL/6JRccHsd, Envigo) were used throughout the study. Viruses were purchased from Penn Vector Core (AAV2/5.CaMKII.GCaMP6f.WPRE.SV40, titer:2.49e13/ml) or custom-made in the case of the retrograde AAV^74^ (retroAAV.EF1a.GCaMP6f.WPRE, Georg Keller, FMI Vector Core, titer: 3.9e11/ml). rAAV2-retro helper was a gift from Alla Karpova & David Schaffer (Addgene plasmid #81070). retroAAV virus was supplemented with blue non-retrograde polymer microspheres (1:2400, Duke Scientific Corp.) to label injection sites. All animal experiments were performed in accordance with institutional guidelines and permitted by the authorities of the canton Basel-Stadt.

### Surgeries

8-week-old male mice were anesthetized (isoflurane) and placed in a stereotactic frame (Kopf Instruments). 300 nl (retroAAV.EF1a.GCaMP6f, ventral hippocampus, medial prefrontal cortex, nucleus accumbens) or 500 nl (AAV2/5.CamKII.GCaMP6f, BA) virus was injected in the respective target area using a glass pipette connected to a picospritzer during a stereotactic brain surgery under isoflurane anesthesia (1-2%). One week after virus injection the animals underwent a second surgery for gradient refractive index (GRIN) lens implantation. An 800-µm-diameter craniotomy was drilled above the BA. Next, a small track was cut with a 0.7 mm sterile needle through the hole and cortex to aid GRIN lens insertion. A 600-µm-diameter GRIN lens (Inscopix) was then slowly advanced into the basal amygdala (−4.4 mm ventral to the pia surface). The GRIN lens was fixed to the skull using light curable glue (Loctite 4305, Henkel) and the skull sealed with Scotchbond (3M), Vetbond (3M) and dental acrylic (Paladur, Kulzer). A custom made head-bar for animal fixation during miniature microscope mounting was attached. Animals were provided with analgesia (buprenorphine, ropivacaine) and their well-being was monitored throughout the experimental period.

### Behavioral paradigm

Animals were single-housed on a 12-hour light cycle and behavioral experiments were performed during the light period. Anxiety tests and fear conditioning were combined in the following manner on seven consecutive days: open field test, elevated plus maze test, habituation 1, habituation 2, fear conditioning, extinction 1, extinction 2. The open field test was performed in a 40 x 40 x 40 cm plastic box with an evenly distributed light intensity of 24 lux. The elevated plus maze was composed of two orthogonal open and closed arms (30 cm) and a center zone (6 cm) elevated at 55 cm above the ground. Mice were allowed to freely explore the open field and elevated plus maze for 10 to 15 min. Habituation and extinction experiments were performed in a 23-cm-diameter circular plexiglass arena. Mice were presented with 5 CS+ and CS−(6 or 12 kHz, counterbalanced) during the habituation sessions and 4 CS− and 12 CS+ during the extinction session. CSs comprised 27 tone pips, each pip 200 ms in duration and 75 dB in amplitude, presented at a rate of 1.1 Hz (Tucker Davis Technologies, TDT 78). Fear conditioning was performed in a 26-cm-wide square plexiglass context using a Coulbourn shock grid. After termination of the CS+ the mouse received a 2-s-long, 0.65 mA AC foot shock 1.1 s after the last tone pip. Behavioral experiments were performed and recorded using Sort Client (2.7.0), Radiant (2.0.0) and Cineplex (3.4.1) software in combination with the CinePlex and MAP data acquisition systems (all Plexon). Animals were perfused with 4% paraformaldehyde (PFA) in phosphate buffered saline (PBS) at the end of the behavioral experiment and brain slices (150 m thick) were cut (vibratome, Leica) and visualized (confocal microscopy, Zeiss LSM700) to verify GRIN lens location and imaging sites. Animals that lacked detectable GCaMP6f expression before baseplate mounting, or had off-target viral injections were excluded from analysis.

### Imaging experiments

Ca^2+^-imaging studies in freely moving mice were performed via the implanted GRIN lens using a miniaturized microscope (nVista, V2, nVista HD 2.0.4 Inscopix)^75^. Microscope base plates were glued to the dental acrylic circa one week before the experiment and mice were habituated to the microscope attachment procedure before behavioral experiments. Miniature microscopes were mounted onto the mouse’s head right before each behavioral experiment using a custom mounting station. Images were acquired at 1024×1024 pixels and at a frame rate of 20 Hz. Imaging parameters were set at 20-80% LED intensity (0.4 −1.7 mW, 473 nm) and a gain of 1-2 depending on GCaMP6f expression levels.

### Data analysis

#### Behavior

Behavioral data was manually scored (EPM, head dips, defined as brief periods of distinct downwards movement of the mouse’s head over the edge of the open arms; OF, rearing) or automatically tracked based on top-down movies using Cineplex Studio and Editor software (Plexon). Freezing was calculated using the 2D motion data and Freezing Analysis plug-in of Editor (Plexon, minimum absence of movement: 2 s, threshold adjusted on a case-by-case basis) combined with additional manual post hoc checks for non-freezing but stationary behaviors (e.g. grooming). Open field test and elevated plus maze 2D location was tracked offline. Open field locations were defined as follows: center: center quarter of open field area, corners: 1/16^th^ of open field area measured from each corner of the arena. Corner entry and exit times were detected as the animal’s crossings of the corner area outlines (36 ± 7 transitions / animal) with a minimum time of 0.5 s in each area before and after transition to exclude events which were non-directed outline crossings (e.g. lingering on the zone border). The number of corner transitions, EPM closed and open arm as well as center location times were calculated using Editor (Plexon) and Matlab.

#### Ca^2+^ imaging

Imaging raw movies were pre-processed and normalized using a fast Fourier transform band pass filter in ImageJ. For normalization, each image was divided by its filtered image. After pre-processing, the movies were spatially down-sampled by a factor of four. The movies of all individual experimental days were then concatenated and motion corrected across all frames in Matlab using the TurboReg algorithm^76^. Single cell ROIs were extracted using a combination of principal and independent component analysis with post-hoc independent component truncation at 50% peak intensity^28,77^. ROI size was limited to a maximum diameter of 30 pxl (ca. 60 µm). ROIs were then overlaid with a maximum intensity projection of the raw movie and excluded, if they overlapped with non-cellular components (e.g. edge of GRIN lens, blood vessels) or if multiple ROIs were detected for the same maximum intensity projection of the same neuron. Pixels within ROIs were normalized and cut at 50% of the maximum ROI pixel intensity. Fluorescence traces for each ROI were extracted as average pixel intensity within the normalized ROI projected along the filtered and motion corrected 20 Hz raw fluorescence movie. Traces which failed to pass quality criteria upon visual inspection across sessions were excluded (i.e. dominated by sharp, negative transients). We typically retained 44% of ROIs (initial ICs: 300) after applying above mentioned exclusion criteria.

Prior to further analysis linear trends across an entire session were removed from the Ca^2+^ traces and z-scores of the detrended traces were used as the activity traces for all further calculations.

#### Area scores

The area bias in single neuron responses was calculated using the average spatial activity map of each neuron: the total activity in a specific x-y location normalized by the total time the animal spent in that location. X-y data was discretized in 256 pixels and smoothed with a 2-d Gaussian kernel that was 1/64 the arena size. The difference between the total normalized activation in the corners (defined as above) and the center was used as a measure of the bias. We calculated the area score for all neurons in the dataset, and set threshold at 1 standard deviation around zero, to capture the behavior of the distribution tails. This threshold was calculated once for the entire population of neurons and applied to each mouse. To create the temporally shifted dataset, behavioral time courses were circle-shifted by a random amount relative to the neuronal activity time course, and area scores were calculated for 1000 random shifts for each animal.

#### Area decoding

We used Support Vector classification with a quadratic kernel for al decoders. Animal location was determined in 1-s bins as center/corners/walls according to the definitions above, and bins with non-consistent behavior were excluded from analysis. To avoid very unbalanced designs, if an animal spent less than 15% of the session in the corner, the decoder was trained and tested on the times the mouse was either in the corner or in the center. Likewise, if the animal spent less than 15% of the session in the center, animal location was defined as corner vs. non-corner locations (center/walls) and the decoder was trained and tested on this distinction. To control for the local dependencies in both the behavioral data and the neuronal responses, we repeated the decoding training procedure with the behavior circularly rotated relative to the neuronal activity. Maintaining the signal’s local structure resulted in a consistent modest drop in decoding performance, that was highly significant (decoding performance on real data was higher than performance on shifted data in 99.8 ± 0.2% of all possible shifts see Fig. S2L).

#### Neuronal response analysis and clustering

Single neuron responses to corner entry were averaged for each neuron in a time window starting 2 s before transition to 5 s after. The mean responses of all neurons from each animal were then clustered into 3 groups using k-means clustering with Euclidean distance between the mean response traces as the distance measure. The result was a cluster of generally activated (ensemble 1), generally inhibited (ensemble 2) and cells with weak / mixed responses (neutral cluster) for each animal. The same procedure was used to define three functional ensembles based on the mean responses aligned to the freezing start period (Fig. 5D).

CS and US responses were analyzed using a combined statistical and supervised cluster analysis. First, CS and US responsive cells were identified as significantly responsive if their binned Ca^2+^ fluorescence (CS: 1 s bins, ±30 s window around CS onset; US 1 s bin, ±14 s window around US onset) during the stimulus was significantly increased (CS) as well as significantly increased or decreased (US), Wilcoxon Signed Rank test, alpha-level: 0.01) compared to baseline conditions in at least 3 or 2 stimulus presentations for the US or CS, respectively (on at least one experimental condition for CS-responses). This minimum number of sensory responses allowed a reliable detection of CS and US plasticity profiles instead of merely general responsiveness across all days, , without being too sensitive for random Ca^2+^ responses during individual tone presentations. Responses for CS were collected to tones 1-5 in the two habituation sessions and in the first extinction session (Ext. early), and to tones 8-12 in the second fear extinction session (Ext. late). The average neuronal responses across both habituation sessions was used as the habituation response of the cells. Next we used a supervised clustering approach on the subset of significantly responsive cells to identify different subtypes of CS and US responsive neurons. Neuronal responses were collected in time windows of [−2 s 15 s] and [−10 s 30 s] around CS and US onset respectively. Principal component analysis was performed on the concatenated responses to the 5 US in the FC session to identify the dynamics in the US response within this session. Responses were then projected on the first 4 principle components (> 60% variance explained) and K-means clustering performed (k = 11, cosine distance). We then manually joined clusters with similar response profiles in relation to the 6 response types described in the article (Fig. 5 and Fig. S6). This procedure was replicated for the CS responses after averaging responses to the 5 selected tones in the habituation and extinction sessions as described above, to identify the different dynamics in CS responses along the conditioning paradigm.

#### Correlation analysis

Average responses aligned to the onset of different behaviors were calculated in the different sessions: corner entry (OF), corner exit (OF), freezing period start and end (habituation, extinction), and head dips (EPM). Response to freezing and corner entry/exit was quantified as the difference between the mean response in the 2 s time window starting 1 s after behavior onset (response) and the 2s time window ending 1 s before onset (baseline). Head dip was quantified as the difference between the mean response in the 1s time window starting 0.8s before behavior detection (response) and the 1 s time window ending 1 s before behavior detection (baseline), due the different dynamics of the behavior. CS and US responses were quantified with a similar baseline period defined as the 2 s time window ending 1 s before stimulus onset and a response period define as the 2 sec after onset (US) or the 9 s starting 1 s after onset (CS). Pearson’s correlation was calculated between the neuronal responses to each pair of behaviors for each animal.

### Statistics

Statistical analysis was performed with Prism 7 (GraphPad) and Matlab (Mathworks). Values are represented as mean ± s.e.m. unless stated otherwise. Box and whisker plots indicate median, interquartile range and the minimum to maximum values of the data distribution. Normality of the data was not assumed and all tests were non-parametric tests. Statistical tests and test statics are mentioned in the text and figure legends. *, ** and *** indicate p-values smaller than 0.05, 0.01 and 0.001, respectively. All data and code will be available upon reasonable request.

---

**Fig. S1.**
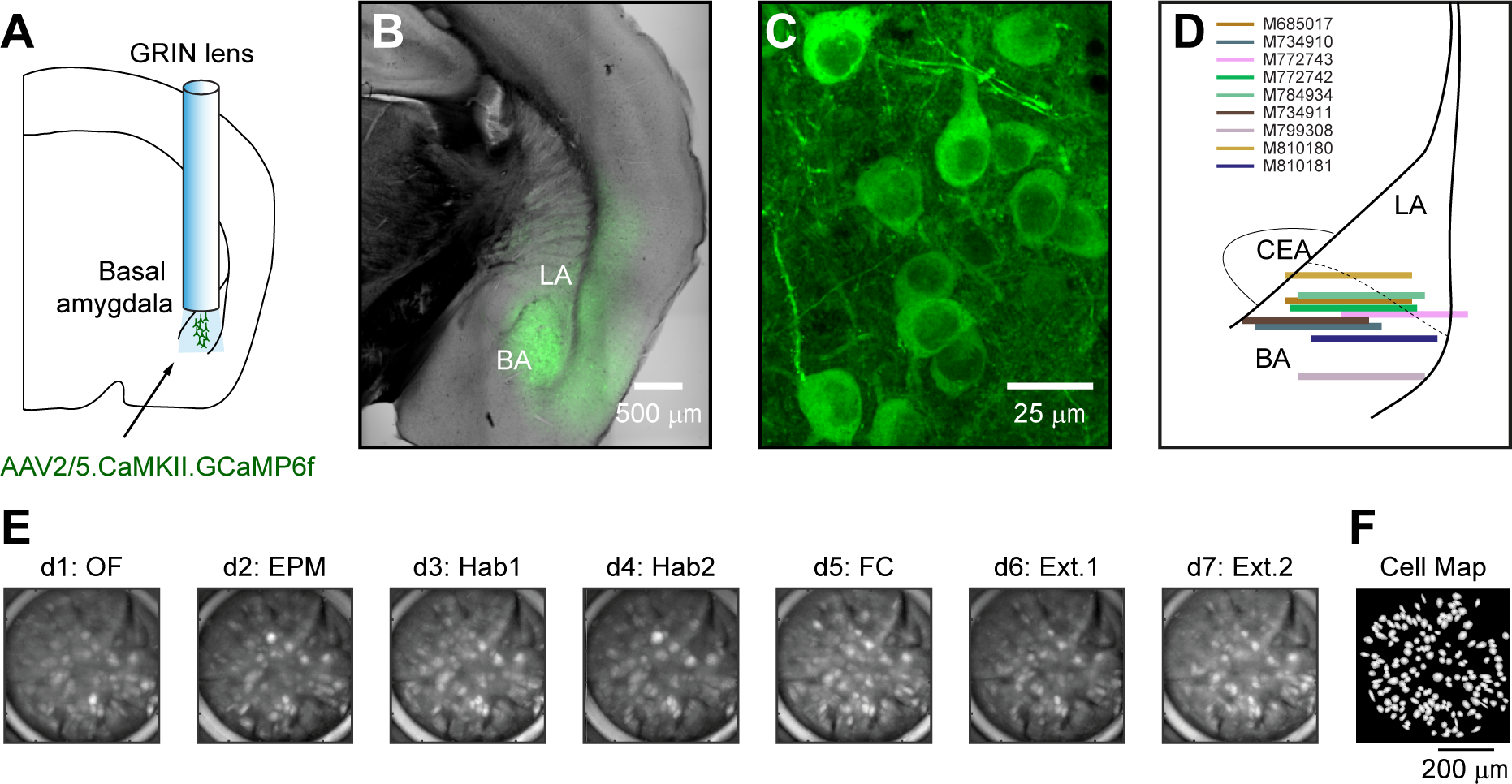
Deep brain imaging of basal amygdala neurons in freely moving animals. **A**) Anatomical scheme of virus injection and GRIN lens location. **B**) Anatomical overview (transmission PMT) of GCaMP6f expression (green) in basal amygdala neurons. **C**) Individual basal amygdala neurons expressing GCaMP6f (maximum intensity projection). **D**) GRIN lens front location of animals expressing GCaMP6f non-specifically in large populations of basal amygdala neurons under the control of the CaMKII promotor (N = 9 mice). **E**) Aligned miniature microscope field of view across seven recording days. OF: open field test. EPM: elevated plus maze test. Hab1/2: habituation day 1/2. FC: fear conditioning day. Ext. 1/2: Extinction training day 1/2. **F**) Map of identified cell regions of interest (ROI). ROIs were identified based on the aligned concatenated imaging movies across all experimental days.

**Fig. S2.**
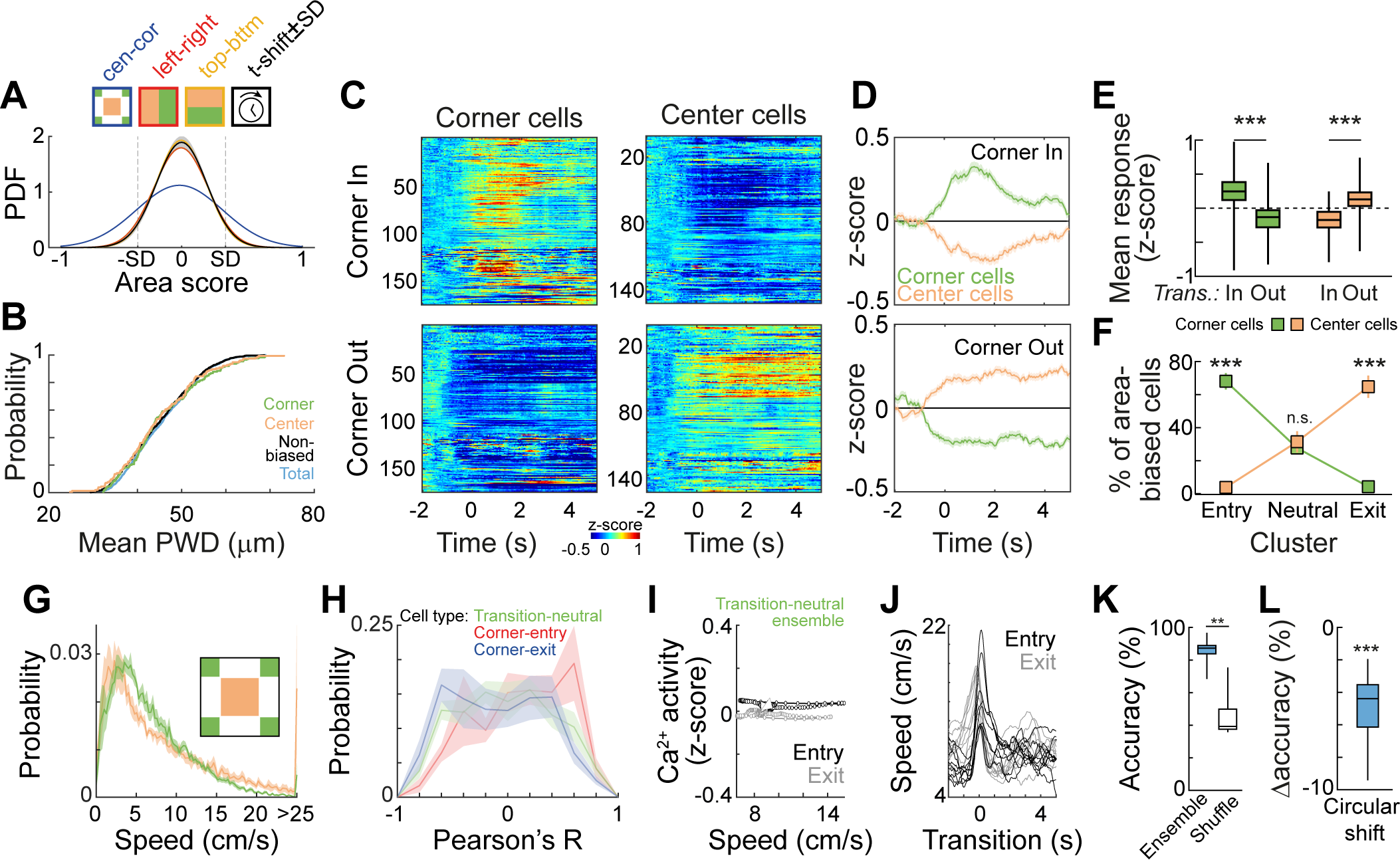
Corner and center cell location, responses upon area transition and speed correlations. **A**) Probability density function (Gaussian fit) of area scores based on differential activity in centers vs. corners (cen-cor, blue) as well as neutral division of the open field (left-right, red; top-bottom, yellow, n = 1201 cells out of N = 9 animals, plotted SD = standard deviation of cor-cen, SD(cor-cen) = 0.36, SD(left-right) = 0.22, SD(top-bottom) = 0.21, Brown-Forsythe Test: F = 177.4, P = <0.001) or temporally shifted data (black ± SD, 100 random circular temporal shifts, SD(t-shift) = 0.21 ± 0.01, all shifts had lower SDs compared to SD(cor-cen)). **B**) Cumulative probability distribution of the mean cellular pairwise distance of all corner cells, center cells, non-modulated cells and the total cell population reveals that the three cell groups are not spatially clustered (Corner vs. total: P = 0.299, Center vs. total: P = 0.414, Non-biased vs. total: P = 0.276, Kolmogorov-Smirnov test). **C**) Average response of all corner and center cells upon in corner and out corner transitions. **D**) Average corner (green) and center (orange) cell responses to in corner (top) and out corner (bottom) transitions (mean ± s.e.m of all corner and center cells). **E**) Average corner and center cell responses to transitions in and out of the corner (Corner cell: In = 0.25 ± 0.02 z-score, Out: −0.16 ± 0.02 z-score, P<0.0001, N = 174 cells; Center cell: In = −0.18 ± 0.01 z-score, Out: 0.15 ± 0.01 zS, P < 0.0001, N = 155, Wilcoxon matched-pairs signed rank test). **F**) Distribution of corner and center cells across different clusters based on in corner transitions (N = 9 animals, Corner entry cluster: Corner vs. Center cells: P < 0.001, Transition neutral cluster: Corner vs. Center cells: P = 0.927, Corner exit cluster: Corner vs. Center cells: P < 0.001, Test: two-way ANOVA with Sidak’s multiple comparisons post hoc test). **G**) Distribution of animal speed in the corners and center of the open field arena (mean ± s.e.m of the distributions of N = 9 animals). **H**) Distribution of individual cell speed Pearson’s correlations for Corner-entry, Corner-exit and Transition-neutral ensembles (mean ± s.e.m of the distributions of N = 9 animals). **I**) Ca^2+^ activity in relation to corner entry (black) and corner exit (grey) transition speed for the transition-neutral ensembles (see Fig. 2G; average response profile for N = 9 animals). Triangles mark the start of the transition. **J**) Mean speed profile upon corner entry (black) and corner exit (grey) for all 9 animals. **K**) Spatial decoding accuracy of corner and center location in actual and time-shuffled data (P = 0.004, Wilcoxon matched-pairs signed rank test). **L**) Average change in decoder accuracy (−5.0 ± 0.7 %) upon circular temporal shift of neuronal activity (N = 9 animals, P = 0.001, One sample T-Test). Box and whisker plots indicate median, interquartile range and the minimum to maximum values of the data distribution.

**Fig. S3.**
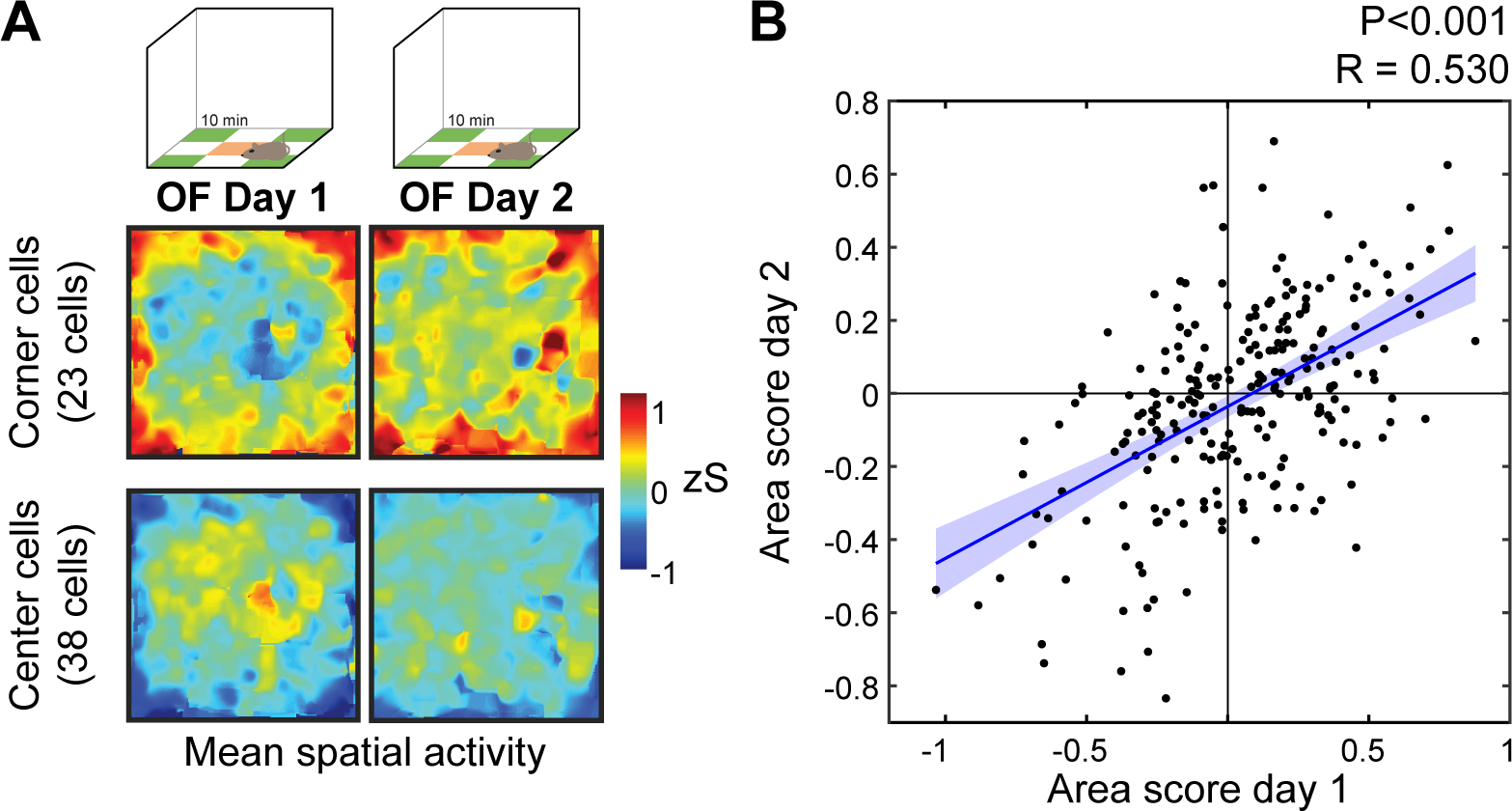
Area coding is stable upon OF re-exposure. **A)** Two day open field (OF) re-exposure paradigm. Average cell activity maps of corner (top) and center cells (bottom) on day 1 (left) and day 2 (right) of open field exposure. **B**) Area score correlation of all recorded cells (area coding and non-coding cells, black dots) on day 1 and day 2 (n = 240 cells from N = 3 mice). Line: linear fit with 95 % confidence bounds. 79 % of identified corner-and center cells displayed stable area score membership across days (48 out of 61 area coding cells).

**Fig. S4.**
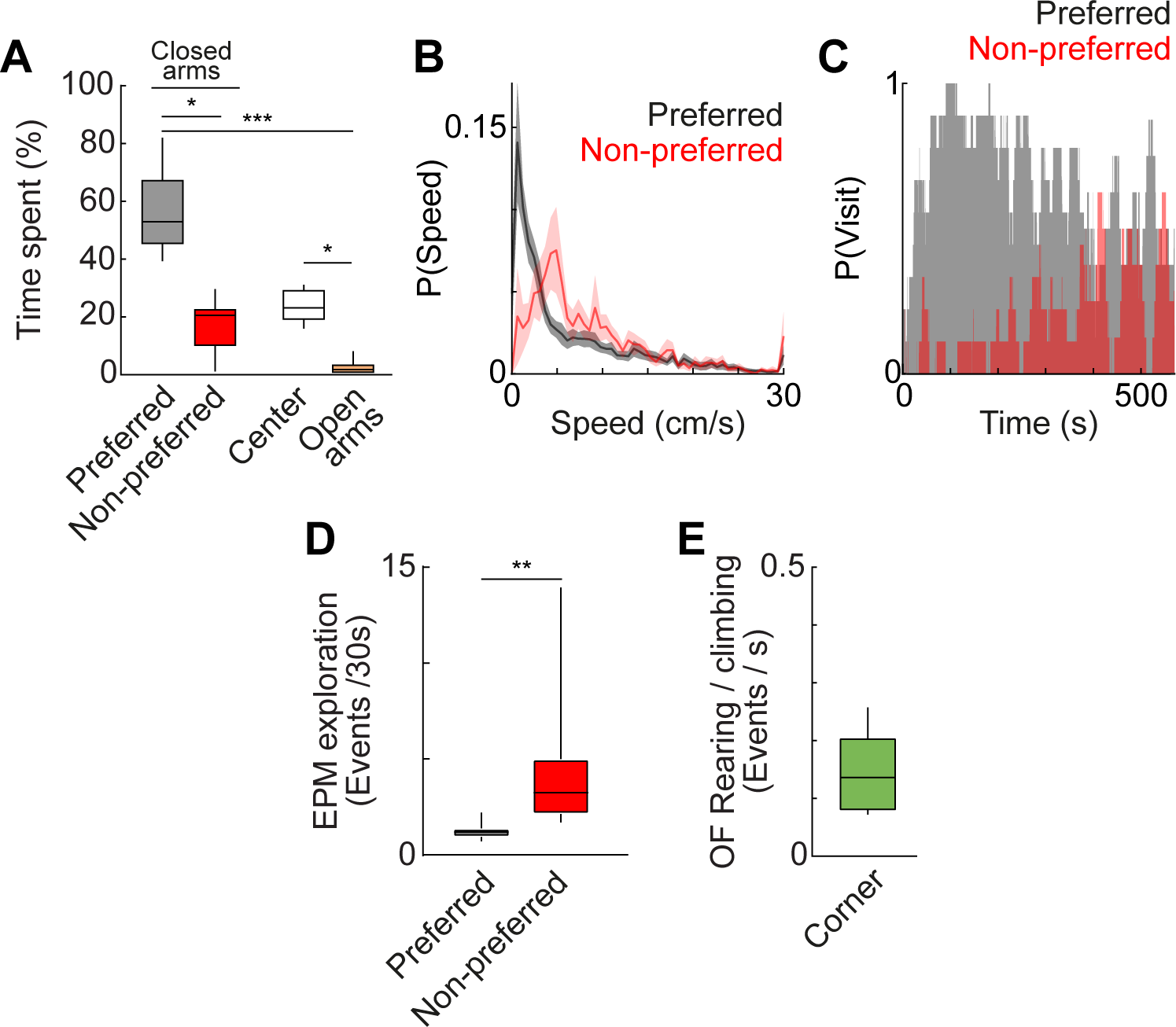
EPM and open field behavior. **A)** Average time spent in the preferred (57 ± 5 %) and non-preferred (17 ± 3 %) closed arms as well as center (24 ± 2 %) of the EPM and its open (3 ± 1 %) arms. Statistically significant p-values: preferred closed vs. non-preferred closed: 0.012, preferred closed vs. open: P < 0.001, center vs. open: 0.012. Friedman test with *post hoc* Dunn’s multiple comparisons test. **B**) Speed distribution in the preferred and non-preferred closed arms of the EPM indicates that the animals are much more exploratory in the non-preferred closed arm. **C**) Probability of preferred and non-preferred closed arm visits throughout the EPM experiment, indicating stability of arm preference. **D**) Exploratory event (rearing and stretch attend posture) frequency in the preferred (1.23 ± 0.14 / 30s) and non-preferred (4.35 ± 1.27 / 30s) closed arm (P = 0.0039, Wilcoxon matched-pairs signed rank test). **E**) Average time-normalized rearing and climbing events in OF corners (0.14 ± 0.02 events/s). N = 9 animals. Box and whisker plots indicate median, interquartile range and the minimum to maximum values of the data distribution.

**Fig. S5.**
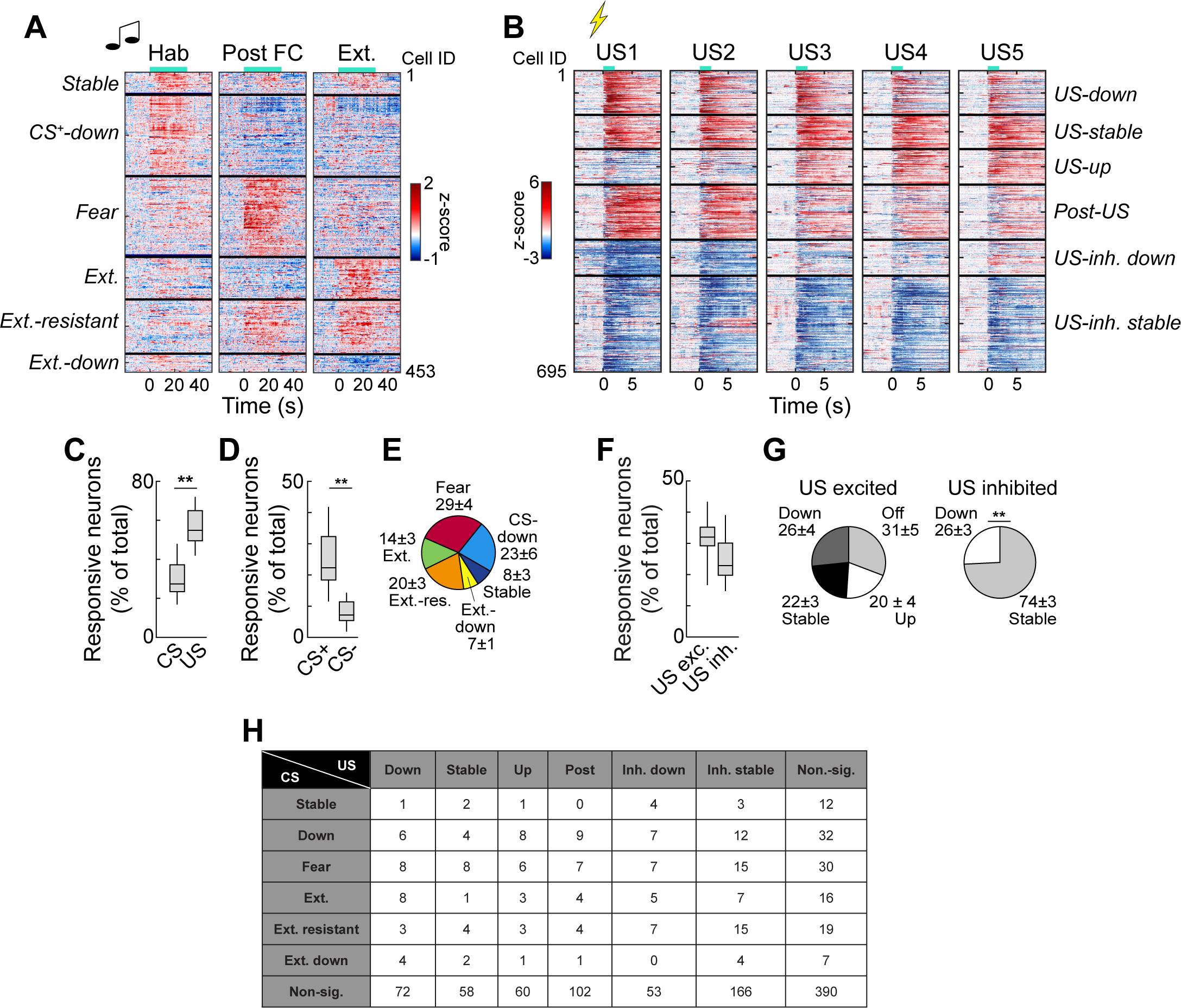
Clusters of CS and US responses. **A)** Average CS+ Ca^2+^ signal of all tone-responsive basal amygdala neurons on three different experimental days. Left: Average response of 10 CS+ responses during habituation day 1 and 2. Middle: Average of the first 5 CS+ responses post fear conditioning on extinction day 1. Right: Average of the last 5 CS+ responses after fear extinction on extinction day 2 (n = 453 neurons from 9 animals). **B**) US Ca^2+^ signal of all shock-responsive basolateral amygdala neurons during fear conditioning. US1-5 indicate the five consecutive USs during tone-shock pairings (n = 695 neurons from 9 mice). Horizontal black lines indicate the cluster borders between the different plasticity types of tone-or shock-responsive neurons, respectively. Horizontal turquois lines indicate start of the CS (tone pips) or US, respectively. **C**) Proportion of CS and US responsive neurons across animals. **D**) Proportion of CS+ and CS-responsive neurons. **E**) Proportion of subgroups of CS plastic neurons. **F**) Proportion of US excited and US inhibited neurons across animals (N = 9 animals, P = 0.129, Test = Wilcoxon matched-pairs signed rank test). **G**) Proportion of subgroups of US plastic neurons. **H**) Total number of intersected functional classes of CS+ and US responsive neurons (data from 9 mice, P = 0.4336, Chi-square test). Box and whisker plots indicate median, interquartile range and the minimum to maximum values of the data distribution.

**Fig. S6.**
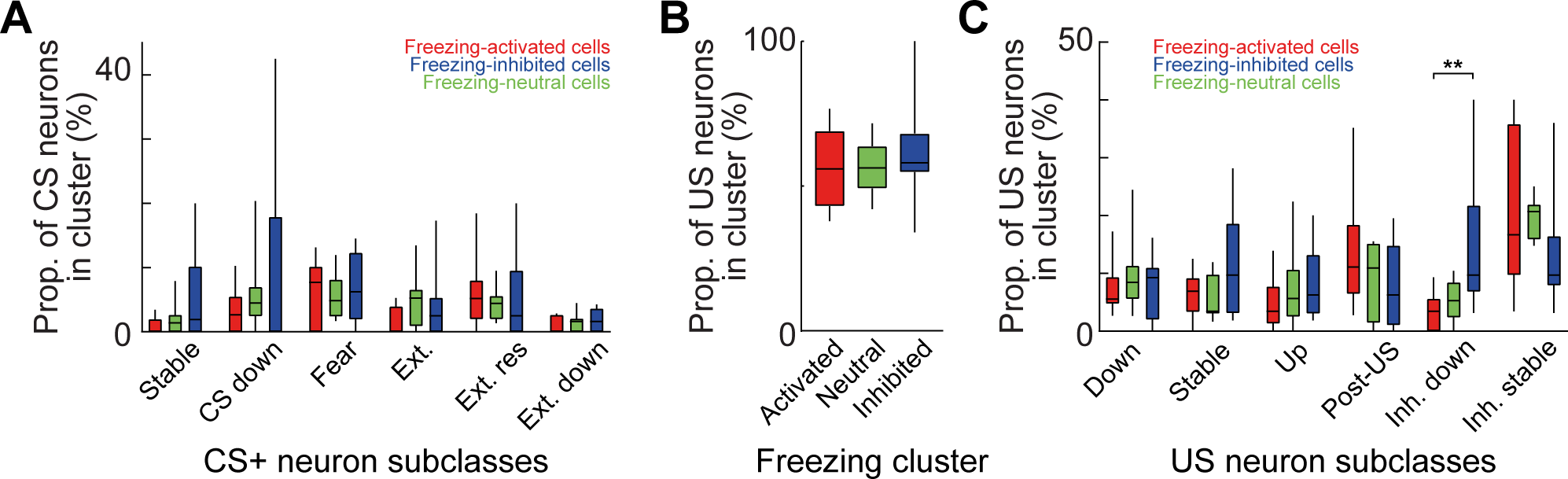
Overlap of CS / US and freezing coding subgroups. **A)** Average proportion of CS+ coding subgroups in freezing activated (cluster 1), freezing neutral and freezing inhibited neurons (cluster 2). No significant difference in the proportion of CS+ coding subgroups in the three freezing subgroups could be found (Stable: P = 0.532, CS down: P = 0.547, Fear: P > 0.999, Ext.: P = 0.475, Ext. resistant: P = 0.685, Ext. down: P = 0.784, Friedman test). **B**) Proportion of general US-responsive neurons in freezing-up, neutral and down-regulated neurons (P = 0.187, Friedman test). **C**) Average proportion of US coding subgroups in freezing activated (cluster 1), freezing neutral and freezing inhibited neurons (cluster 2). No significant difference in the proportion of US coding subgroups in the three freezing subgroups could be found (Down: P = 0.569, Stable: P = 0.569, Up: P = 0.654, Post-US: P = 0.741, Inh. down: P = 0.007, Dunn’s multiple comparisons test: Cluster 1 vs. Cluster 2: P = 0.0096, Inh.: P = 0.057, all Friedman test). Box and whisker plots indicate median, interquartile range and the minimum to maximum values of the data distribution. All data is generated from N = 9 animals.

**Fig. S7.**
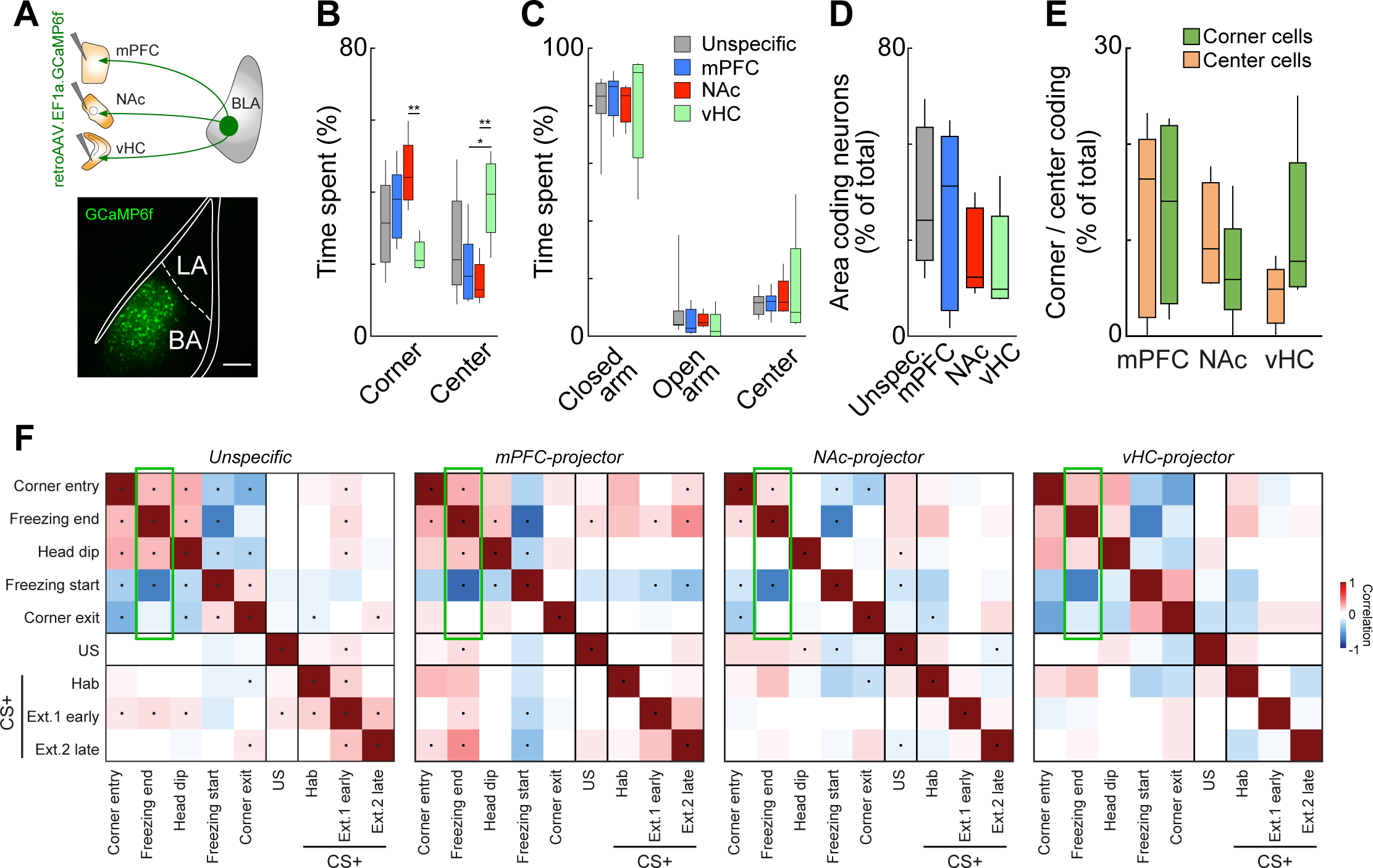
Area specific coding and neuronal population correlations across behavior and sensory representations of defined basal amygdala projection neurons. **A)** Viral strategy for retrograde AAV labelling of amygdala projection neurons with GCaMP6f. GCaMP6f expression in mPFC-projecting BA neurons. Scale bar: 200 μm. **B**) Summary data of open field behavior in unspecific and projection neuron groups. Corner NAc vs. Corner vHC: P = 0.0071, Corner mPFC vs. Corner vHC: P = 0.017, Corner NAc vs. Corner vHC: P = 0.005, Two-way ANOVA, Tukey multiple comparison. **C**) Summary data of elevated plus maze behavior for unspecific and projection neuron groups. Two-way ANOVA, Tukey multiple comparison. No difference between projector groups. **D**) Summary data of area-biased neurons for all groups. P = 0.255, Kruskal-Wallis test. **E**) Summary of corner and center cell proportions in projection neurons. No significant difference between groups. Wilcoxon matched-pairs signed rank test. **F**) Correlation matrices of population responses during exploratory and freezing behavior as well as CS and US responses for unspecific neurons, mPFC-projectors, NAc-projectors and vHC-projectors. Green rectangles indicate state coding correlations. Box and whisker plots indicate median, interquartile range and the minimum to maximum values of the data distribution. All data is generated from the following number of animals: unspecific: 9, mPFC: 6, NAc: 5, vHC: 5 (except panel f: vHC = 4).

**Fig. S8.**
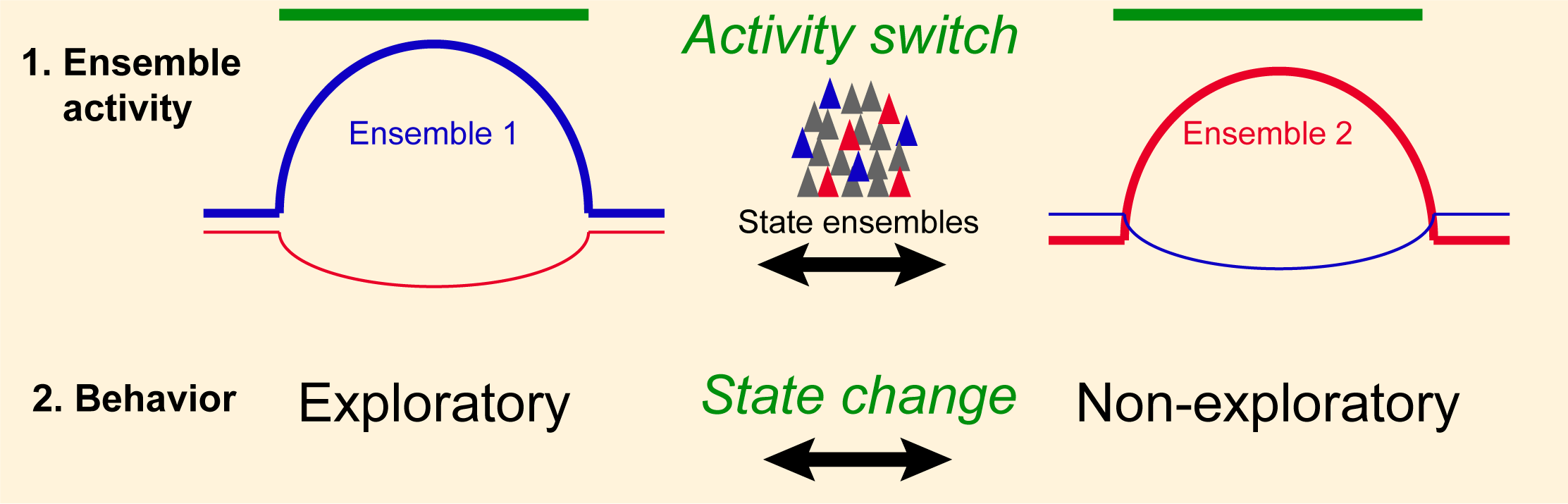
Two large, non-overlapping amygdala populations encode behavioral states. State changes from exploratory (e.g. corner behavior, exploration in the open field, head dips, reduction of freezing, reduced anxiety) to non-exploratory behaviors (e.g. center behavior, homebase arm retreat, start of freezing, anxiety-like, defensive) are encoded by two, opposingly activated, non-overlapping ensembles of basal amygdala principal neurons. This ensemble activity signature is recurring across different behaviors associated with the same state, and switches sign when the animal enters an antagonistic state.

**Fig. S9.**
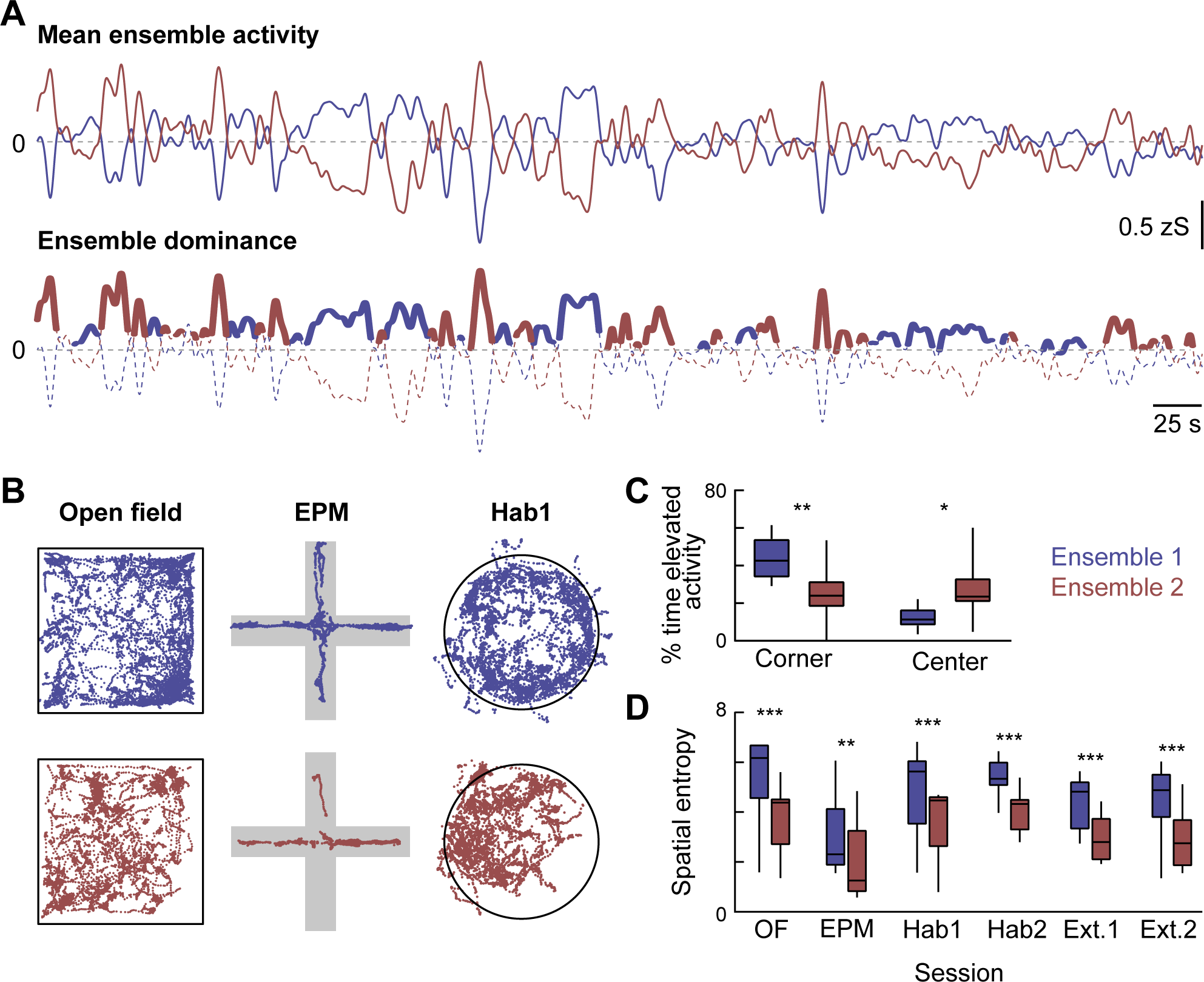
Amygdala ensembles dynamics predict behavioral states. **A)** Top: Average signal of two main ensembles in the basal amygdala. Ensemble 1/2 cell identity was based on k-means clustering for two clusters (correlation distance) of the neuronal responses in the first 5 minutes of the open field session and fixed across all subsequent behavioral paradigms. Bottom: Episodes of specific elevated activity in relation to the opposing ensemble (cut-off: 70^th^ percentile of difference signal). **B**) Behavioral track during episodes of elevated activity in ensemble 1 (top row) and ensemble 2 (bottom row) for the open field (left), elevated plus maze (middle) and habituation day 1 (right). Tracks outside the arena of Hab1 illustrate rearing events. **C**) Fraction (%) of time of elevated ensemble activity the animals were in the corner or center of the open field. Two-way ANOVA with Sidak’s multiple comparisons test: F(1,32) = 13.5, P < 0.001. **D**) Spatial entropy based on XY location data during specific elevated activity in ensemble 1 or ensemble 2. N = 9 animals. Two-way ANOVA with Sidak’s multiple comparisons test: F(1,8) = 163.9, P < 0.0001. Box and whisker plots indicate median, interquartile range and the minimum to maximum values of the data distribution.

